# Diversity and biogeography of SAR11 bacteria from the Arctic Ocean

**DOI:** 10.1101/517433

**Authors:** Susanne Kraemer, Arthi Ramachandran, David Colatriano, Connie Lovejoy, David A. Walsh

## Abstract

The Arctic Ocean is relatively isolated from other oceans and consists of strongly stratified water masses with distinct histories, nutrient, temperature and salinity characteristics, therefore providing an optimal environment to investigate local adaptation. The globally distributed SAR11 bacterial group consists of multiple ecotypes that are associated with particular marine environments, yet relatively little is known about Arctic SAR11 diversity. Here, we examined SAR11 diversity using ITS analysis and metagenome-assembled genomes (MAGs). Arctic SAR11 assemblages were comprised of the S1a, S1b, S2, and S3 clades, and structured by water mass and depth. The fresher surface layer was dominated by an ecotype (S3-derived P3.2) previously associated with Arctic and brackish water. In contrast, deeper waters of Pacific origin were dominated by the P2.3 ecotype of the S2 clade, within which we identified a novel subdivision (P2.3s1) that was rare outside the Arctic Ocean. Arctic S2-derived SAR11 MAGs were restricted to high latitudes and included MAGs related to the recently defined S2b subclade, a finding consistent with bi-polar ecotypes and Arctic endemism. These results place the stratified Arctic Ocean into the SAR11 global biogeography and have identified SAR11 lineages for future investigation of adaptive evolution in the Arctic Ocean.

## Introduction

The SAR11 (Pelagibacterales) group accounts for roughly 30% of bacteria in the ocean surface and 25% of mesopelagic bacteria [1–3]. High phylogenetic diversity and divergence into ecological lineages (i.e. ecotypes) of SAR11 tends to mirror distinct conditions in the oceanic environment [2, 4–6]. SAR11 are classified into clades and subclades based on 16S rRNA genes and further classified into phylotypes based on rRNA internal transcribed spacers (ITS) (Table 1). Three major SAR11 clades (S1, S2 and S3) are currently recognized, with several subclades defined within S1 (S1a, S1b, S1c). Within this diversity approximately 12 phylotypes have been described [2, 5]. To date, many of the phylotypes are thought to have restricted distributions. For example, there are three phylotypes in subclade S1a: P1a.1 is associated with cold environments [2], while P1a.2 and P1a.3 are associated with temperate and tropical environments, respectively [2, 5]. Subclade S1b is generally associated with tropical environments [2, 5, 7], while subclade S1c is associated with deep marine samples [5]. Recently, the clade S2 was divided into subclades S2a and S2b. The S2a subclade is further divided into an oxygen minimum zone subclade (S2a.a) and a tropical subclade (S2a.b) [7]. The S2 subclade phylotypes have been further divided by environment. P2.1, is associated with tropical environments [2], P2.2 is reported from cold, especially Antarctic, waters [2, 8], and the P2.3 phylotype is more ambiguous; associated with both cold waters and tropical deep-sea environments [2, 5]. To date, three phylotypes have been distinguished for clade S3; P3.1 is associated with coastal, mesohaline surface [9] and tropical samples [2], P3.2 is found in brackish [9], temperate [2], as well as Arctic samples [10] and the third S3 phylotype, LD12, is found in freshwater [11] (Table 1).

**Table 1:**
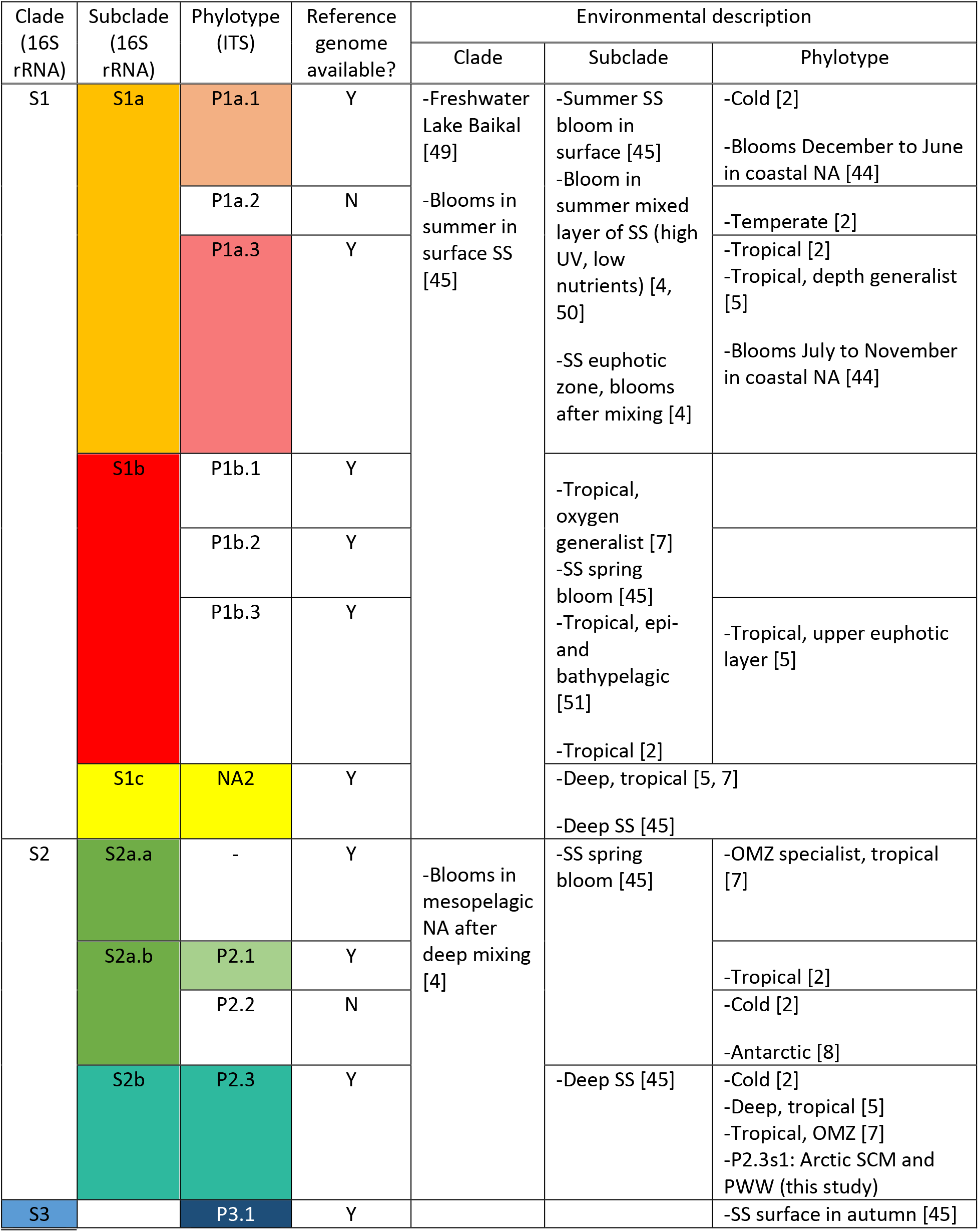
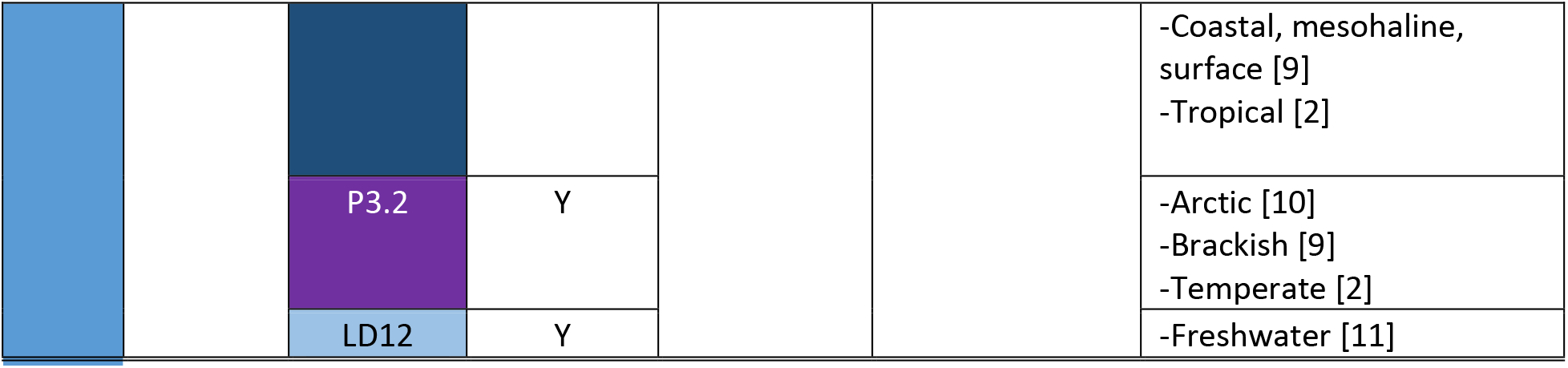
SAR11 clades, subclades and phylotypes, as well as their environmental associations as described in the literature (SS: Sargasso Sea).

The Arctic Ocean is small, geographically isolated, and more influenced by freshwater and ice compared to other oceans [12–14]. These distinct characteristics would favour the evolution of locally adapted microbial assemblages, however evidence for this is difficult to document. Biogeographic patterns of taxa reported from the Arctic vary, with reports of cosmopolitan, bi-polar and endemic species, suggesting a potential for different levels of specialization to local conditions [6, 15–17]. Although SAR11 are reported widely and account for 25 to 30% of Arctic Ocean bacterial assemblages [18, 19], comparatively little information exists on Arctic SAR11 diversity [2]. The extensive knowledge of SAR11 diversity and ecology in other oceans makes it an attractive clade to investigate the potential for ecotypes adapted to Arctic Ocean conditions. The objective of this study was to place SAR11 from the Arctic Ocean within a global context using ITS phylogenetic analysis and comparative genomics using metagenome-assembled genomes (MAGs). We then examined the distribution of SAR11 along a latitudinal transect of the stratified waters of the Canada Basin in the western Arctic Ocean. We targeted three water masses in the upper 200 m, the surface layer, the deep chlorophyll maximum (DCM), which corresponds to a halocline formed by Pacific Summer Water, and the Pacific Winter Water (PWW) layer [20]. These three water masses were previously found to have distinct microbial communities [21] and we hypothesised that 1) different SAR11 ecotypes would be favored within them, and 2) that the distinct characteristics of the Arctic Ocean would favor the existence of endemic SAR11 ecotypes

## Methods

### Sampling and metagenomic data generation

Samples were collected aboard the Canadian Coast Guard Icebreaker CCGS Louis S. St-Laurent from the Western Arctic Ocean from latitudes 73° to 79° N in October 2015, during the Joint Ocean Ice Study cruise in the Canada Basin (Table 2). Sample collection and preservation, DNA extraction, and metagenomic data generation were as described previously [22], and further details given in the Supplementary Information. The metagenomic data is deposited in the Integrated Microbial Genomes database at the Joint Genome Institute at https://img.jgi.doe.gov, GOLD project ID Ga0133547.

**Table 2:** Western Arctic Ocean metagenome sampling sites, their environmental feature and environmental parameters including sampling depth (Depth), water temperature (Temp.), Salinity, Chl a Fluorescence, CDOM, Oxygen and Nitrate (NO_3_) limit of detection 0.02 (nd)

| Station | Lat. | Long. | Feature | Depth [m] | Temp. [°C] | Salinity | Fluorescence [mg m<sup>-3</sup>] | CDOM [mg m<sup>-3</sup>] | Oxygen [mmol m<sup>-3</sup>] | NO<sub>3</sub> [mmol m<sup>-3</sup>] |
| --- | --- | --- | --- | --- | --- | --- | --- | --- | --- | --- |
| CB2_154 | 73.22 | -150.22 | SUR | 7 | -1.26 | 25.70 | 0.15 | 2.54 | 391.85 | nd |
| CB2_152 | 73.22 | -150.22 | DCM | 67 | -0.86 | 31.48 | 0.33 | 3.9 | 351.56 | 4.45 |
| CB2_150 | 73.22 | -150.22 | PWW | 177 | -1.45 | 33.18 | 0.05 | 4.21 | 281.85 | 15.98 |
| CB4_138 | 75.26 | -150.07 | SUR | 5 | -1.39 | 26.14 | 0.12 | 2.58 | 394.08 | nd |
| CB4_136 | 75.26 | -150.07 | DCM | 79 | -0.02 | 31.16 | 0.23 | 3.9 | 329.14 | 4.65 |
| CB4_134 | 75.26 | -150.07 | PWW | 208 | -1.47 | 33.14 | 0.05 | 4.36 | 282.39 | 16.13 |
| CB8_130 | 77.1 | -150.23 | SUR | 5 | -1.46 | 27.2 | 0.19 | 2.78 | 395.60 | nd |
| CB8_128 | 77.1 | -150.23 | DCM | 58 | -0.15 | 30.97 | 0.30 | 3.89 | 383.72 | 0.35 |
| CB8_126 | 77.1 | -150.23 | PWW | 213 | -1.45 | 33.14 | 0.05 | 4.42 | 281.09 | 16.24 |
| CB11_90 | 79.25 | -150.06 | SUR | 5 | -1.48 | 27.31 | 0.24 | 2.67 | 394.88 | nd |
| CB11_88 | 79.25 | -150.06 | DCM | 25 | -1.04 | 29.69 | 0.25 | 3.22 | 401.18 | nd |
| CB11_86 | 79.25 | -150.06 | PWW | 190 | -1.46 | 33.15 | 0.05 | 4.35 | 281.89 | 15.82 |

### ITS phylogenetic analysis

SAR11 ITS sequences were retrieved from twelve assembled metagenomes, clustered and filtered (Supplementary information). We combined the Arctic ITS sequences with reference sequences from SAR11 genomes and ITS sequences from two previous biogeographic studies [2, 5] and assigned Arctic ITS sequences to phylotypes. We determined phylotype distribution across the samples with PCoA ordination of the Bray-Curtis dissimilarity as described in the Supplementary Information.

### Metagenome-assembled genomes

We conducted metagenomic binning using the Metabat2 pipeline [23] on a 12 sample coassembly [22, 24]. Further SAR11 bin filtering and cleaning was conducted as described in the Supplementary Information. Seven SAR11 MAGs with low contamination values (<4%) and relatively high completeness values estimated using CheckM [25] (36 % to 47%) were selected for further analysis (Table 3) (Genbank accession numbers VBTQ00000000, VBTR00000000, VBTS00000000, VBTT00000000, VBTU00000000, VBTV00000000, VBTW00000000). The distribution of orthologs across SAR11 MAGs and reference genomes was analyzed with ProteinOrtho [26]. A set of 39 single copy ortholog genes were concatenated and used for phylogenetic analysis using MEGA-cc with a JTT substitution model [27], as further described in the Supplementary Information.

**Table 3:** Summary statistics of MAGs from the Western Arctic Ocean.

| **Subclade** | **MAG** | **Size (mb)** | **Cov (x)** | **GC (%)** | **Completeness (%)** | **Contamination (%)** | **N50 (kb)** | **# of scaffolds** | **# of genes** | **# of unique genes** |
| --- | --- | --- | --- | --- | --- | --- | --- | --- | --- | --- |
| S1a | 312 | 0.42 | 27.83 | 27.3 | 47.17 | 1.89 | 21.23 | 20 | 462 | 33 |
| S2a | 112 | 0.41 | 8.66 | 28.31 | 38.68 | 1.89 | 18.76 | 20 | 462 | 49 |
| S2a | 272 | 0.3 | 17.96 | 29.52 | 35.85 | 0 | 98.98 | 5 | 320 | 10 |
| S2a | 410 | 0.42 | 16.77 | 30.68 | 24.53 | 1.89 | 18.96 | 26 | 462 | 36 |
| S2a | 484 | 0.44 | 38.18 | 28.54 | 35.09 | 0 | 38.17 | 14 | 481 | 35 |
| S2 | 144 | 0.63 | 27.78 | 29.21 | 41.51 | 3.77 | 30.34 | 23 | 672 | 47 |
| S2 | 196 | 0.39 | 23.43 | 28.5 | 46.99 | 0 | 93.24 | 8 | 430 | 29 |

### Comparative genome content, average nucleotide identity and signatures of selection of Arctic MAGs

To identify Arctic-specific SAR11 genes, we compared all genes within the Arctic MAGs to those found within 41 SAR11 reference genomes using ProteinOrtho [26]. The protein functions of genes only found in Arctic MAGs were retrieved following the IMG annotation [28] or the SwissProt database [29]. We calculated average nucleotide identity (ANI) between Arctic MAGs and reference genomes representing the different subgroups following the method implemented within IMG [28].

### Fragment recruitment and SAR11 biogeography

To determine the prevalence of Arctic MAGs across global marine biomes, we performed reciprocal best hit analysis [22, 30] for 163 metagenomic samples from a range of marine biomes. Metagenomic datasets included 129 TARA ocean datasets [31] which were randomly sub-sampled to a size of approximately 1 GB of reads each to facilitate analysis. These were added to the Arctic metagenomes (also randomly subsetted to 1.5 GB of reads) and 22 Antarctic datasets that included 20 marine and 2 ACE lake samples [2]. We examined the distribution of the seven Arctic MAGs and 48 representative SAR11 reference genomes as described in the Supplemental Information. Best hits from the reciprocal blast were filtered to a minimum length of 100 bp and a minimum identity of 98% and the number of recorded hits per reference genome and metagenomic sample were used to calculate the number of reads recruited per kilobase genome per gigabase metagenome (RPKG). See Supplemental Table 1 for all metagenomic datasets and reference genomes.

We utilized the RPKG matrix for PCoA ordination of its Bray-Curtis dissimilarity. The envdist function as implemented in vegan with 999 permutations was used for *Post-Hoc* tests of environmental variables. [32].

## Results

### Environmental setting

The stratified upper waters of the Canada Basin in the Western Arctic Ocean sampled during the late summer–autumn of 2015 were typical for the region, with warmer and fresher summer waters above colder slightly saltier winter Pacific-origin water (Table 2). Salinity at the surface ranged from 25.6-27.3 and nitrate concentrations were below the detection limit (0.5 μM). Below the surface water, relative chlorophyll *a* fluorescence at the DCM was calculated to range from 0.23 to 0.33 mg m^−3^. The DCM depth varied from 25-79 m, with the shallowest value at CB11, the most northerly station along the western edge of the Canada Basin (Figure 1A-B). Nitrate concentrations increased with depth to up to 16 mg m^−3^ in the PWW, where chlorophyll *a* was low (0.05 µmol m^−3^).

**Figure 1:**
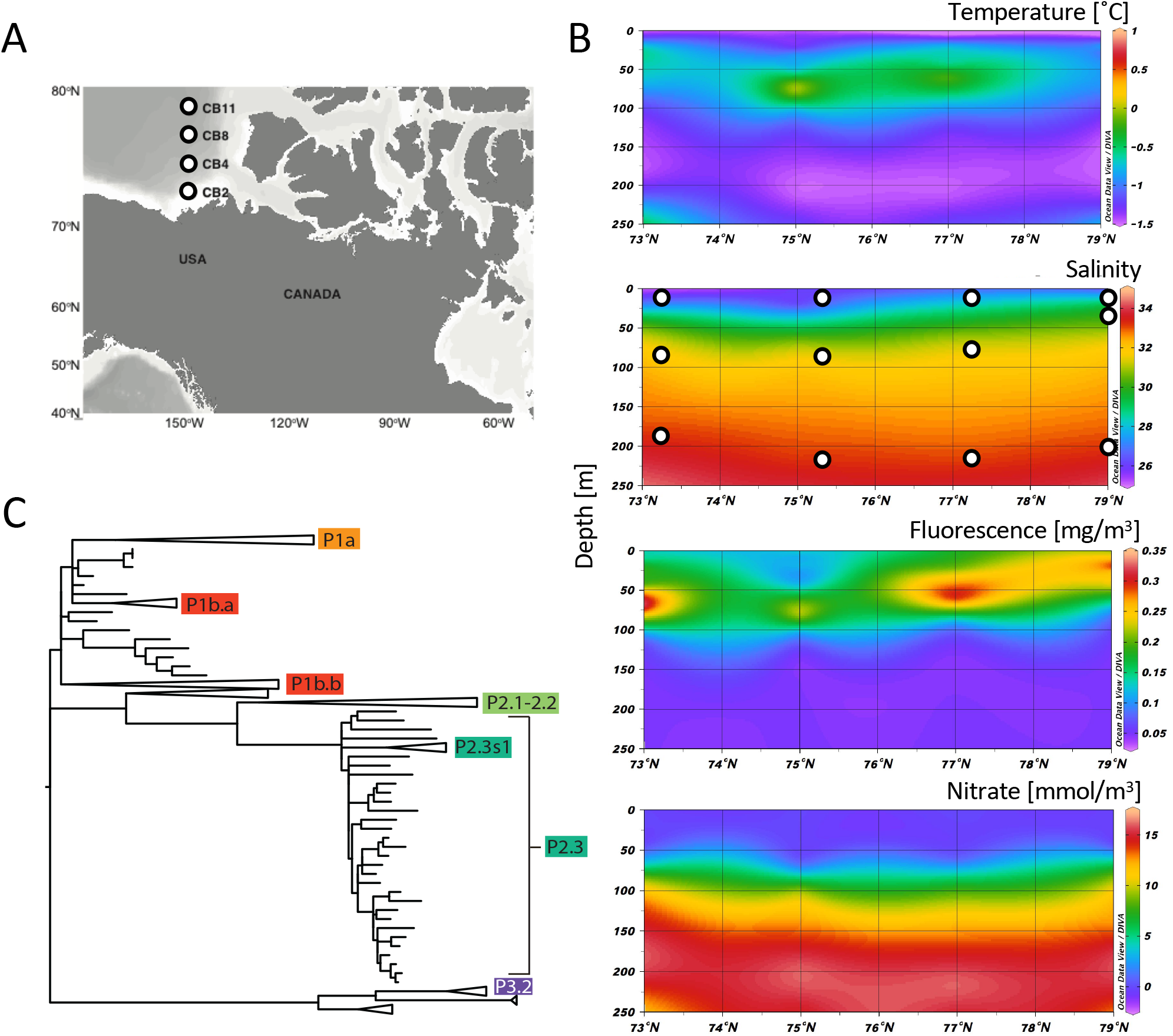
Study metadata and overview of SAR11 ITS phylogeny. A) Sampling locations of the Western Arctic Ocean metagenomes. B) Environmental profiles of sampling locations showing temperature (°C), salinity, chlorophyll *a* fluorescence (mg m^−3^) and nitrate (mmol m^−3^). C) Maximum likelihood ITS phylogeny including Arctic and reference sequences. Only subclades which contain Arctic ITS sequence types are labeled.

### SAR11 ITS sequence diversity

No SAR11 16S rRNA sequences were present in the metagenomes. Within the metagenomic assemblies, we identified 140 high quality SAR11 ITS sequences which were clustered into 111 unique ITS sequence variants (SVs) and combined these SVs with previously published sequences for phylogenetic analysis. Sixty Arctic ITS SVs did not cover the full ITS region and were assigned to phylotypes using their best BLAST hit against full-length SAR11 ITS SVs. In total, 6 distinct phylotypes were evident from ITS SVs (Figure 1C), from all major clades: S1, S2, and S3 (Figure 1C, Supplemental Figure 1A-D).

#### Clade S1

Within S1, we identified the S1a and S1b subclades. The Arctic ITS S1a SVs mostly clustered apart from previously published P1a SVs (Supplemental Figure 1A). These P1a-related SVs were common in samples from the PWW and DCM layer, but nearly absent from the surface layer (Supplemental Figure 1A, Figure 2A). Within subclade S1b (Supplemental Figure 1B), Arctic Ocean ITS SVs were found in two clusters. One cluster corresponded to a previously described P1b.a group [2], while the other grouped within a previously designated non-monophyletic tropical P1b cluster [2], hereafter termed P1b.b (Supplemental Figure 1B), but we were unable to connect these groupings to the three previously described P1b.1-3 phylotypes [5]. P1b.a contained three full-length Arctic SVs and was found nearly exclusively in the PWW (Supplemental Figure 1B, Figure 2A).

**Figure 2:**
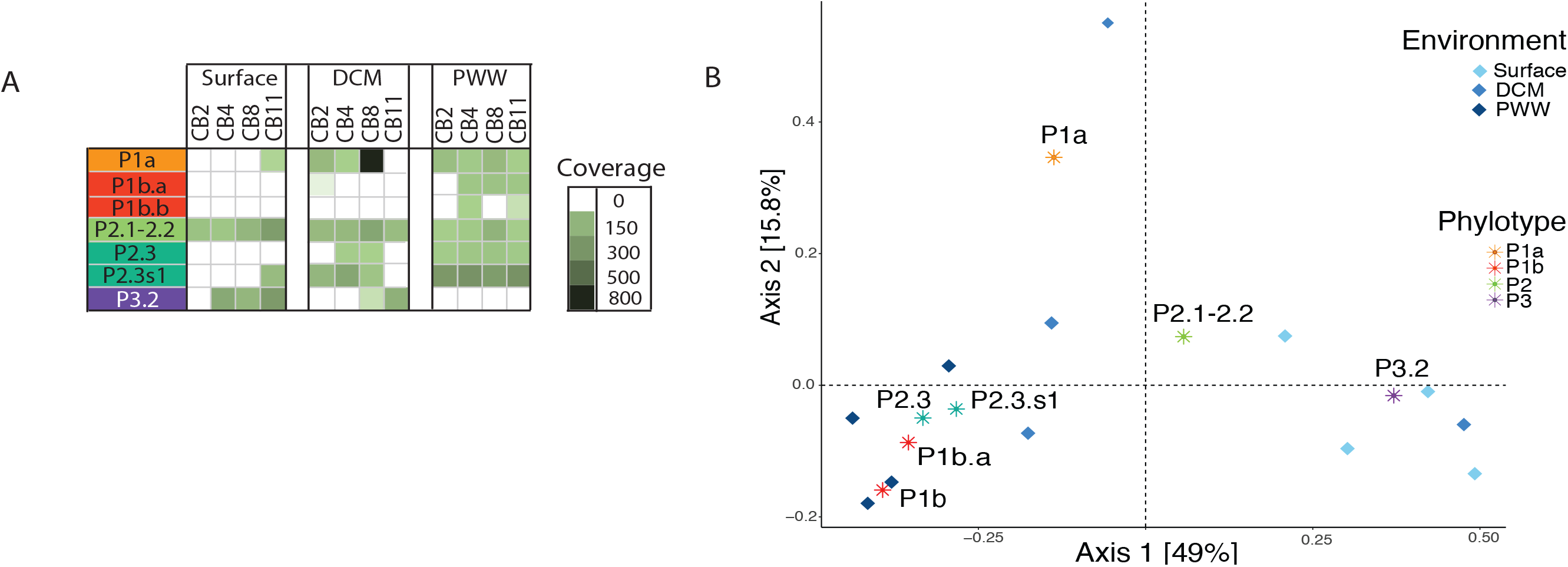
Distribution of SAR11 phylotypes in the Arctic Ocean. A) ITS region coverage of Arctic SAR11 phylotypes across samples. B) Principal coordinate analysis ordination of Bray-Curtis dissimilarities of Arctic samples based on the coverage of different ITS phylotypes. Scaling 2 is shown. Dark blue diamonds: PWW samples, blue diamonds: SCM samples, light blue samples: surface samples. Asterisks show phylotypes’ weighted average frequency.

#### Clade S2

The majority of ITS SVs were assigned to phylotypes within subclade S2 (Supplemental Figure 1C). In our phylogenetic analysis, we recovered a monophyletic group within the P2.3 phylotype, with Arctic SVs clustering with two deep water Red Sea SVs, forming a novel cluster P2.3s1 (Figure 1C, Supplemental Figure 1C). Apart from a single DCM sample, which contained a high number of P1a sequences (coverage of ~670, compared to an average coverage of ~40, Fig. 2A), SVs within the P2.3s1 phylotype were the most frequently detected in Arctic waters, specifically in DCM and PWW samples (Supplemental Figure 1C, Figure 2A). The published ITS SVs previously assigned to P2.1 and P2.2 were interspersed with one another, and we therefore refer to this subgroup as P2.1–2.2. P2.1–2.2 was distributed relatively evenly across sampling depths and locations (Supplemental Figure 1C, Figure 2A).

#### Clade S3

We recovered two ITS SVs belonging to the brackish Arctic phylotype P3.2 (Supplemental Figure 1D). P3.2 was common in the less saline surface water samples and absent below the DCM (Supplemental Figure 1D, Figure 2A).

### Phylotype abundance and biogeography

PCoA ordination of samples based on the relative abundance of phylotypes showed that SAR11 assemblages were structured along the first axis in relation to the water layer sampled (Figure 2B). The second axis had less explanatory power and assemblage structuring was mostly driven by the highly abundant P1a SV in the DCM sample of station CB8. PWW samples contained a high contribution of diverse phylotypes including P1a, P1b.a, P1b, P2.1–P2.2 and P2.3 and P2.3s1, while P3.2 was most frequent in surface layer samples. P2.1–P2.2, was not associated with contributions to a specific depth class of samples likely because multiple poorly resolved ecotypes are contained within this group (Figure 2A, B). Secondary fitting of environmental variables onto the ordination did not yield significant results.

### Characteristics of Arctic Ocean SAR11 MAGs

We binned SAR11 scaffolds from the Arctic Ocean metagenome co-assembly based on tetranucleotide frequency and coverage across samples. After automated binning and manual curation, we selected seven high quality SAR11 MAGs for further analysis (Table 3). Quality was based on high N50 values (19-100Kb), a low number of scaffolds (5-26), relatively high completeness (35-47%), and low contamination (0-4%) values (except for ‘completeness’ following [33]).

While SAR11 ITS sequences were abundant in the metagenomic dataset, none were present in the SAR11 MAGs. Moreover, SAR11 MAGs did not contain rRNA genes and thus their placement into 16S rRNA or ITS-based phylogenies was not possible. Instead we investigated their phylogeny using a concatenated gene tree of Arctic MAGs, reference genomes, and SAGs from the Red Sea and the Eastern Tropical North Pacific oxygen minimum zone (ETNP OMZ) [7] (Figure 3A). The Arctic MAG SAR11–312 belonged to subclade S1a and was prevalent at and below the DCM layer, but not at the surface (Figure 3B). This MAG was more closely related to phylotype P1a.1 than to P1a.3; a placement supported by its maximum ANI value in comparison to a P1a.1 reference genome (80%). Its phylotype could not be determined, as no closely related reference genome contained ITS sequences.

**Figure 3:**
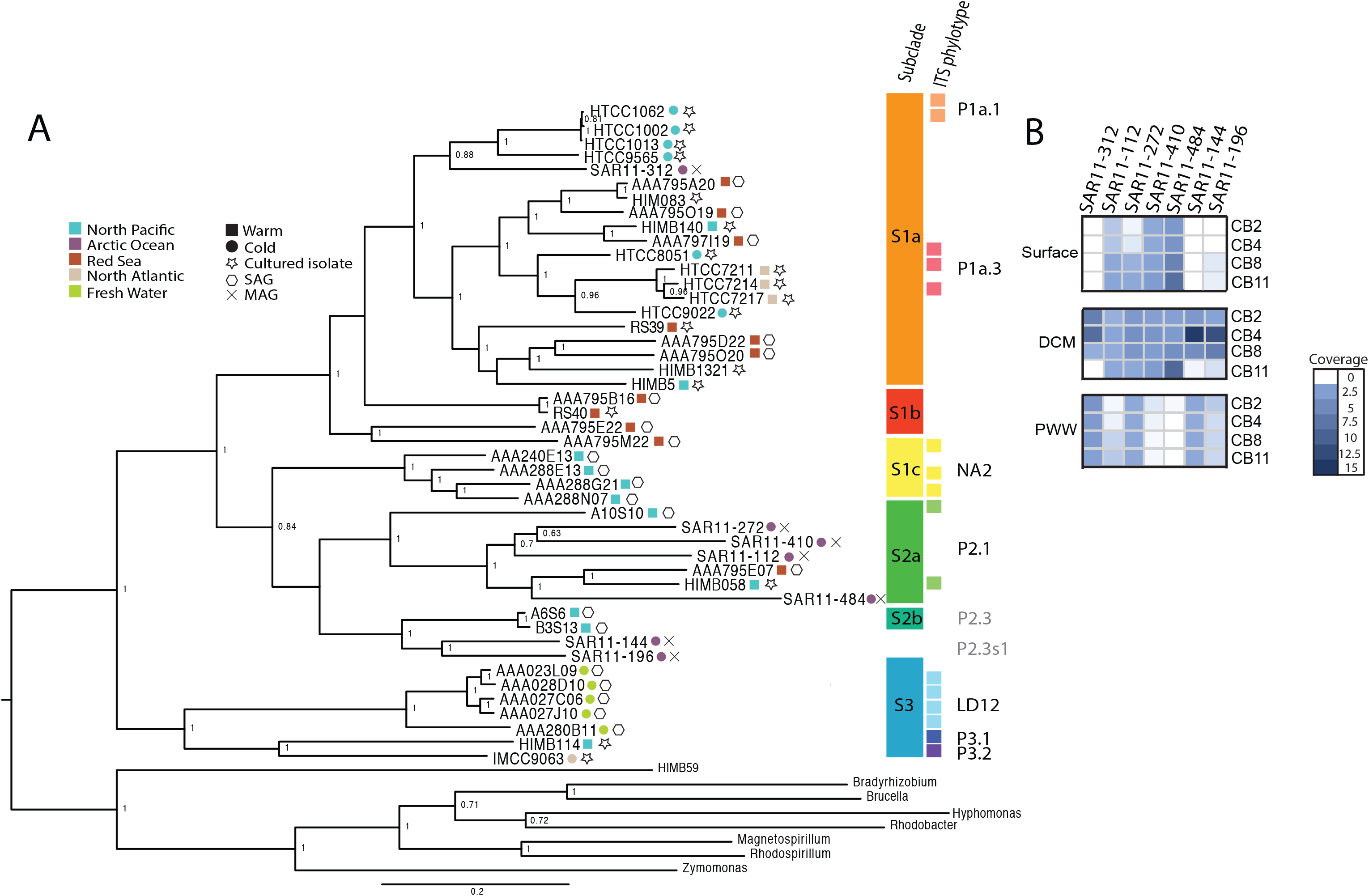
Phylogenetic context and distribution of Arctic SAR11 MAGs. A) Maximum likelihood phylogeny based on 39 concatenated orthologous loci. Only bootstrap values higher than 0.6 are shown on the tree. Colored squares and circles at the tips indicate the environment and temperature of origin (if known), matching phylotypes if known are indicated. Colored squares indicate reference genomes with known ITS phylotypes. Grey labeling indicates inferred phylotypes based on phylogenetic placement and distribution of the genomes. B) Coverage of Arctic MAG scaffolds across Western Arctic ocean metagenomes.

The remaining six MAGs were members of the S2 clade of SAR11. Four of the MAGs (SAR11–112, SAR11–272, SAR11–410, and SAR11-484) were members of the S2a subclade, while two (SAR11–144 and SAR11–196) were most closely related to two SAGs from the S2b subclade that originated from the ETNP OMZ (Figure 3A). The S2a MAGs exhibited differential distributions in the stratified waters of the Arctic Ocean. Three closely related MAGs (SAR11– 112, SAR11–272, SAR11–410) were detected at all depths, while a more distantly related S2a MAG (SAR11–484) was only detected in the surface and DCM samples (Figure 3B). The phylogenetic placement of these MAGs was supported by their maximum ANI values in comparison to the S2a reference genome HIMB058 (>75%). In contrast, MAGs SAR11–144 and SAR11–196 were detected in DCM and PWW samples (Figure 3B) and are the most likely MAGs to represent the novel P2.3s1 phylotype defined in this study. However, as S2b reference genomes lack ITS sequences, we were unable to directly confirm that subclade S2b MAGs belong to the P2.3 phylotype. Due to the absence of any S2b reference genomes with high completeness, we were unable to calculate ANI of SAR11–144 and SAR11–196 with S2b.

### Comparison of gene content

To investigate the potential of local adaptation through variation in gene content we identified SAR11 genes specific to the Arctic Ocean by comparing the distribution of orthologs between Arctic MAGs and 60 reference genomes representing the known phylogenetic diversity of SAR11. Of the 2 648 orthologs we identified across SAR11 genomes, 233 were only found in Arctic MAGs and four of these were present in more than one MAG. The majority of these orthologs were poorly characterized proteins (58%, 136 orthologs: COG categories S and R). A further 24% (55 orthologs) were involved in metabolism, 3% (7 orthologs) in cellular processes and signaling and 7% (16 orthologs) in information storage and processing (Supplemental Table 2).

### Global biogeography of SAR11 genomes

We investigated the biogeographic distribution of Arctic SAR11 populations using Arctic MAGs and available SAR11 genomes. Arctic MAGs were either bi-polarly distributed or only found in the Arctic Ocean. MAG SAR11–312, belonging to subclade S1a, was detected only in the DCM and PWW layer of the Arctic ocean (Figure 4A). Arctic S2a MAGs fell into two groups. SAR11-410, −112 and −484 were found largely in the surface and DCM layers of the Arctic and Southern Ocean, whereas SAR11-272 was detected only in the DCM and PWW samples of the Arctic Ocean. The Arctic MAGs most closely related to the S2b subclade also showed distributions suggesting Arctic endemism and were only found below the surface layer of the Arctic Ocean (Figure 4A).

**Figure 4:**
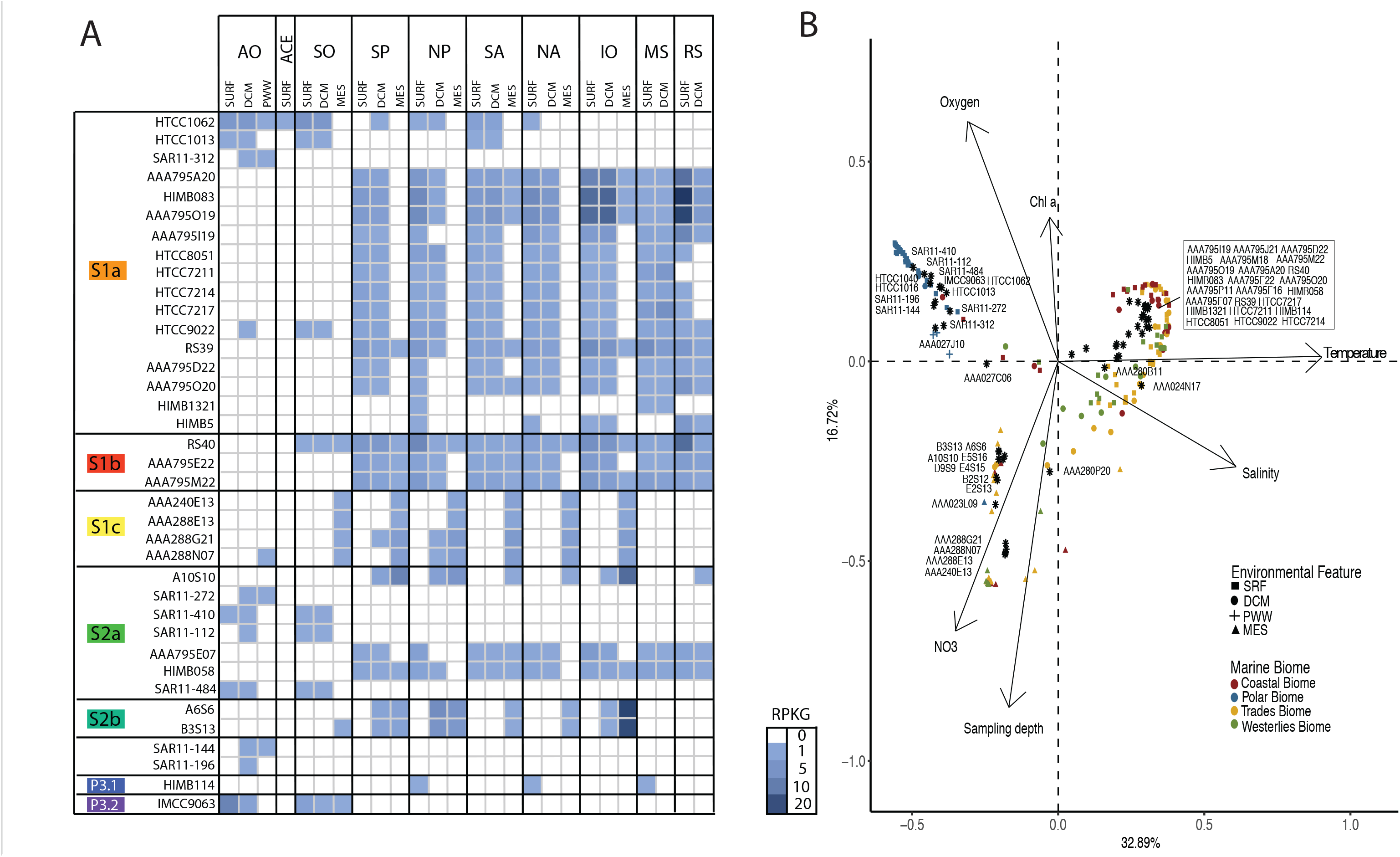
Global biogeography of SAR11 genomes. A) RPKG table of Arctic MAGs and reference genomes by ocean layer and ocean region. Abbreviations for oceans are Arctic (AO), Southern (SO), South Pacific (SP), North Pacific (NP), South Atlantic (SA), North Atlantic (NA), Indian Ocean (IO), Mediterranean (MS) and Red (RS) Seas. For clarity only RPKG values ≥ 0.1 are shown in the figure. We also performed fragment recruitment for five LD12 genomes, but observed no significant recruitment across all metagenomic datasets. B) Principal coordinate analysis ordination of Bray-Curtis dissimilarities of metagenome samples based on RPKG values of arctic MAGs and reference genomes. Scaling 2 is shown. Arrows indicate significant environmental variables after *Post-Hoc* testing. Asterisks show the weighted average of each genomes’ frequency.

Other than the Arctic MAGs, most SAR11 genomes recruited poorly across both Arctic and Antarctic metagenomic samples, suggesting they are not significant members of Arctic SAR11 assemblages. However, there were some exceptions: S1a genome HTCC9022, which was among the most widely distributed genomes, was prevalent in the Antarctic and to a lesser degree the Arctic. Two other S1a genomes, HTCC1062 and HTCC1013, were also prevalent in polar samples. Within the S2 clade, only Arctic MAGs recruited in either Arctic or Antarctic samples, indicating that distinct phylotypes inhabit polar niches, in contrast to more temperate generalist S2a genomes such as HIMB058 and AAA795E07 (Figure 4A). Lastly, S3 genome IMCC9063, characterized as a fresher, arctic phylotype [9] was most prevalent in Arctic surface waters, followed by Antarctic samples.

To further explore SAR11 biogeography in relation to environmental conditions of ocean biomes, we performed PCoA ordination of the Bray-Curtis distance matrix of the RPKG-values (Figure 4B). Overall, metagenomic samples clustered according to their biome and sampling depth. Along the first axis, samples separated according to a temperature and salinity gradient, with colder, fresher samples such as the Arctic and Antarctic environments investigated here distinguished from warmer, saltier environments. Along the second axis, metagenomic samples were separated according to their depth. Deep samples were characterized by high nitrate concentrations while more shallow samples had higher oxygen and Chl *a* concentrations. Notably, Arctic and Antarctic surface and Arctic DCM samples clustered together (Figure 4B, as polar biome samples in the upper left quadrant), while Arctic PWW samples were distinct in their compositions (Figure 4B, blue crosses). S1a MAG SAR11–312 belonged to an apparently cold-adapted S1a cluster consisting of HTCC1062, HTCC1013, HTCC1040 and HTCC1016 which had a vastly different distribution compared to most other S1a genomes, which were found largely in warmer coastal and trades biomes (e.g. RS39, HIMB5, HIMB083). Arctic S2a MAGs were associated with polar DCM and surface samples. The Arctic MAGs SAR11–144 and 196 were similarly associated with polar samples, and differentially distributed from the most closely related North Pacific genomes A6S6 and B3S13.

## Discussion

Ribosomal RNA amplicon surveys of the marine environment have documented the occurrence of SAR11 in Arctic marine systems [34–36], but the diversity and biogeography of the different lineages has not been investigated in detail. Here we found evidence for the presence of at least six SAR11 phylotypes in the Arctic. The global success of the SAR11 group across marine environments points to a capacity to adapt to a wide array of environmental conditions. While Arctic and Antarctic Oceans are superficially similar environments with respect to solar radiation, the presence of sea ice and algae blooms following ice melt in spring [17], the Arctic Ocean is surrounded by land and freshwater input from large rivers makes the Arctic Ocean a more estuary-like marine biome compared to the Antarctic. The rivers also bring in terrestrial-derived organic matter which represents a potential substrate and selective force on the bacterial communities [22]. Increasing evidence suggests microbial Arctic endemism in other bacterial groups [22], but this is the first genome-level study to our knowledge within the SAR11 group.

The biogeography of ITS phylotypes offers a hint that local selective forces are at work but the relatively coarse nature of the marker masks potential differentiation at the genome level, such that ITS-based phylotypes may not necessarily discern discrete bacterial populations that are locally adapted. Recent advances in sequencing techniques and genome assembly algorithms have resulted in the utilization of MAGs and SAGs to extend the biogeographies of bacteria beyond the phylogenetic marker level and infer patterns of local adaptation at the genome level [7, 23, 37]. Utilizing this approach, gene content of individual SAR11 genomes has been linked to niche partitioning based on nutrients [38], metabolic adaptation to environmental productivity [39] and oxygen minimum zones [7]. However, this approach may have limitations since assembling SAR11 genomes from metagenomes is notoriously difficult due to their high levels of polymorphism [40]. Moreover, recent work suggests that gene content differences alone may not be sufficient to explain SAR11 biogeographic patterns [41]. Nonetheless, our combined MAG-based and marker gene-based results converged, providing support for the existence of novel arctic ecotypes.

### Structure of Arctic Ocean SAR11 assemblages

Arctic phylotypes belonging to S1a, S2 and S3 showed preferences linked to different water masses [5]. The fresher Arctic surface waters were dominated by a P3.2 phylotype that had been previously associated with Arctic and brackish waters [9, 10]. In contrast, PWW depths were dominated by the cold [2] and deep [5] phylotype P2.3. The majority of all recovered ITS phylotypes from the Arctic Ocean belonged to the P2.3 phylotype, with one highly abundant phylotype (P2.3s1, Figure 1C) that was previously only represented by a few sequences from deep in the Red Sea [5]. The Arctic Ocean may constitute the centre of the P2.3s1 range distribution [42] and its potential for local adaptation as well as its niche requirements needs further exploration. In contrast to P3.2 and P2.3, the P2.1–P2.2 phylotype showed a broader distribution across depths and samples. However, our phylogenetic analyses failed to recover the two distinct previously published phylotypes P2.1 (tropical) and P2.2 (cold) [2].

To further investigate the link between SAR11 diversity and the environment we utilized metagenomic binning to assemble seven Arctic Ocean SAR11 MAGs that belonged to or were closely related to subclades S1a and S2a and S2b and investigated their biogeographic distribution and transcriptional activity in the Arctic Ocean. Three of the four S2a MAGs recovered here were widely distributed across depth classes and most stringently corresponded to the distribution of the P2.1–P2.2 phylotype. The two MAGs most closely related to S2b reference genomes were mostly present in DCM and PWW waters, mirroring the distribution of the P2.3 and P2.3s1 phylotypes. However, in the absence of direct ITS evidence we were unable to test directly that S2b-related Arctic MAGs correspond to the extremely common P2.3s1 ITS phylotype. Surprisingly, we failed to assemble a S3 MAG, which based on the abundance of corresponding ITS sequences, the high fragment recruitment of the S3 reference genome IMCC9063, should have been recoverable. Moreover, we were unable to detect any long (>10 kb) SAR11 scaffolds mapping to S3. The absence of any other strong S3 signal, apart from the presence of ITS sequences, is intriguing and warrants further investigation.

### Placing Arctic SAR11 into the global biogeography

To determine the biogeography of Arctic MAGs, in comparison to reference genomes representing SAR11 diversity, we performed fragment recruitment analysis across 163 metagenomic datasets from different marine biomes, including the Arctic and two Antarctic habitats: the Southern Ocean and Ace Lake, a saline Antarctic lake. In general, Arctic MAGs showed either Arctic-endemic or bi-polar distributions. Several Arctic S2a MAGs were found only in Arctic and Antarctic biomes, indicating a bi-polar distribution of members of this subclade. One Arctic S1a and S2a MAG, as well as two MAGs which were most closely related to subclade S2b were found only in samples from the Arctic Ocean, providing support for the presence of endemic Arctic SAR11. However, due to the low genome completeness of the MAGs and the lack of closely related reference genomes in the case of the putative S2b genomes it was not feasible to link this potential endemism to gene content [7, 39] or selection acting on genes responsible for the apparent local adaptation to the Arctic, as was recently done for a different and wide-spread SAR11 population [41].

It has been discussed whether SAR11 diversity is shaped by neutral evolution [43], or whether the multitude of subclades found within SAR11 represent ecotypes adapted to local environmental conditions (e.g. [44]). Hellweger and colleagues [43] found that distinct populations characterized by up to 0.5% diversity (roughly corresponding, for example, to the rRNA gene diversity between phylotypes P1a.1 and P1a.3 [44]) can arise neutrally. However, a range of other studies [1, 4, 45] have described seasonal and spatial patterning consistent with adaptation to environmental conditions and colonization of environmental niches. Consistent with ecotype adaptations to local conditions, we found that SAR11 assemblages within the metagenomic datasets to be structured by both their biomes and environmental features. This finding, in correspondence with the fact that diversity between most SAR11 subclades and phylotypes is higher than 0.5% [2, 5], indicates that the majority of diversity within the clade is likely to be shaped by adaptation to environmental niches.

The majority of SAR11 reference genomes (including most S1a and S1b genomes), are associated with metagenomic datasets from warmer, more shallow environments from the Trades, Coastal and Westerlies Biomes. A second group of reference genomes (mostly S1c) were most common in deeper water metagenomic datasets with lower oxygen and higher nitrate concentrations, as previously reported for S1c genotypes found in nitrate replete oceanic OMZs [7]. Polar biome metagenomic datasets formed a distinct third environmental group. Within the polar datasets, those from the PWW clustered away from all other environments, providing support for the hypothesis that distinct SAR11 assemblages in the samples are selected by environmental conditions, specifically with respect to the potentially endemic Arctic S1a and S2 MAGs, which were frequent here and in Arctic DCM samples. In contrast to PWW metagenomic datasets, other Arctic datasets clustered closely with those from the Southern Ocean, pointing towards the bi-polar distribution for several of the Arctic S2a MAGs found in these datasets.

Seasonal variability of SAR11 phylotypes in association with mixing and phytoplankton bloom events has been extensively described at the Bermuda Atlantic Time-series Study (BATS) site in the Sargasso Sea [4, 45] as well as at coastal environments [44]. The absence of seasonal data in our study makes a direct inference of Arctic phylotype seasonality impossible, but Arctic summer/fall patterns can be compared to those described in other marine biomes. In contrast to summer samples from the BATS, phylotypes belonging to S1a and S1b were relatively scarce in the Arctic surface layer [4, 45], though P1a was abundant in most Arctic DCM samples., Brackish phylotype P3.2 was abundant in most Arctic surface water samples, which is an analogous distribution to phylotype P3.1 that blooms in BATS surface waters in the fall [45]. Whether the apparently more cold-tolerating P3.2 phylotype fills the same seasonal niche as P3.1 in tropical and temperate waters remains to be investigated. S2 phylotypes have been found year-round in deeper waters, but bloom specifically in the spring within the upper mesopelagic at BATS [4, 45], where they are likely involved in DOM remineralization following winter deep mixing [4]. In contrast, Arctic S2a MAGs were found to be abundant in the euphotic zones of only Arctic and Antarctic marine biomes, indicating that the MAGs’ niche may be spatially, rather than temporally, defined. In agreement with previous work, we found the S2b reference genomes A6S6 and B3S13 to be highly abundant in the DCM and mesopelagic layer in the North Pacific but absent in the Arctic Ocean. The absence of any S2b reference genome hits in the Arctic Ocean could point towards Arctic S2b-like MAGs (and the putatively corresponding P2.3s1 phylotype) replacing them as endemic Arctic specialists in the DCM and PWW, but, in the absence of seasonal samples, further work is needed to elucidate the seasonal patterns of Arctic ecotypes.

Different degrees of endemism within bacterial communities in the Arctic Ocean have been reported. Patterns vary for different groups of bacteria, but in general Arctic bacterial communities are complex assemblages of bacteria with cosmopolitan, bi-polar, and Arctic-specific distributions, indicating varying degrees of adaptation to local conditions. [6, 16, 17, 46– 48]. In the present study, we set out to describe and place Arctic Ocean SAR11 into the global SAR11 biogeography using both marker gene and MAG-based approaches. Assembly of the highly genome-streamlined SAR11 clade, which shows high rates of recombination, from metagenomes is difficult, complicating comparisons with 16S rRNA-based approaches. Nevertheless, we detected a previously nearly undescribed ITS phylotype, P2.3s1, which was the most common phylotype in most Arctic PWW and DCM samples. Moving from phylotypes to MAGs, we detected Arctic SAR11 genomes with both apparently endemic and bi-polar biogeographic distributions. The selective forces shaping these biogeographic distributions as well as the resulting adaptive responses on the genome level merit future investigation.

## Acknowledgements

Data were collected aboard the CCGS Louis S. St-Laurent in collaboration with researchers from Fisheries and Oceans Canada at the Institute of Ocean Sciences and Woods Hole Oceanographic Institution’s Beaufort Gyre Exploration Program and are available at http://www.whoi.edu/beaufortgyre. We would like to thank both the captain and crew of the CCGS Louis S. St-Laurent, the chief scientist, William J. Williams, and the scientific team aboard. The work was conducted in collaboration with the U.S. Department of Energy Joint Genome Institute, a DOE Office of Science User facility, and was supported under Contract No. DE-AC02-05CH11231. Funding Discovery grants (D.W. and C.L.). This study is also a contribution to ArcticNet, a Network of Centers of Excellence (Canada). The Canadian Natural Science and Engineering Research Council (NSERC) Discovery (C.L and D.W) and Northern Supplement (C.L), the Fonds de recherche du Québec Nature et Technologies (FRQNT) supporting Québec-Océan (C.L. D.W) and the Canada Research Chair Program (D.W.) are acknowledged.

## Conflict of interest statement

The authors declare no conflict of interest.

## Supplementary Information

Supplementary methods description (docx).

Supplemental Information: Supplemental Figure Legends (docx).

